# RenSeq and whole genome sequencing uncover allelic diversity of clubroot resistance genes in commercial breeding canola lines

**DOI:** 10.64898/2026.07.22.740042

**Authors:** Jiaxu Wu, Soham Mukhopadhyay, Muhammad Asim Javed, Yanick Asselin, Elisa Fantino, Coreen Franke, Edel Pérez-López

**Affiliations:** Département de Phytologie, Faculté des Sciences de l’agriculture et de l’alimentation, Université Laval, Québec City, QC, Canada, G1V 0A6; Centre de Recherche et d’innovation sur les Végétaux (CRIV), Université Laval, Québec City, QC, Canada, G1V 0A6; Institute de Biologie Intégrative et des Systèmes (IBIS), Université Laval, Québec City, QC, Canada, G1V 0A6; L’Institute EDS, Université Laval, Québec City, QC, Canada, G1V 0A6; Nutrien Ag Solutions Canada, Saskatoon, SK, Canada, S4N 4L8

**Keywords:** Brassica napus, effector-triggered immunity, RenSeq, clubroot resistant gene, NLR annotation

## Abstract

Clubroot disease, caused by the obligate biotrophic pathogen Plasmodiophora brassicae, is a major threat to canola (Brassica napus) production worldwide. Clubroot-resistant (CR) cultivars remain the most effective disease-management strategy, but the genetic basis of resistance in commercial canola remains poorly understood because many resistance sources are proprietary and associated genotypic information is rarely accessible. Although nucleotide-binding leucine-rich repeat (NLR) immune receptors account for most cloned CR genes, no pan-NLRome has incorporated CR lines used in commercial canola breeding. Here, we combined whole-genome sequencing and resistance gene enrichment sequencing (RenSeq) to assemble and annotate the NLR repertoires of five homozygous CR inbred lines (IH1–IH5) used for commercial breeding and displaying contrasting resistance profiles against predominant Canadian P. brassicae pathotypes. We integrated these NLRomes with the susceptible cultivar Westar to construct a comparative pan-NLRome for canola. Across the five CR lines, total NLR content was highly conserved, ranging from 504 to 517 genes, with TIR-NLRs representing the predominant class. C-JID-containing TIR-NLRs accounted for more than 30% of each NLR repertoire, and integrated-domain analysis identified conserved and genotype-specific NLR-IDs, including previously unreported domains in IH4. Pan-NLRome analysis resolved 366 NLR orthogroups (OGs), 60.7% of which were core, and identified resistant-line-enriched OGs absent from Westar as candidate CR-associated loci. Unexpectedly, a homolog of the functionally characterized CR gene, CRa, was detected in five CR lines. Moreover, a homolog of another CR gene, Crr1a, was detected in both resistant and susceptible lines, indicating that the presence/absence of a gene alone does not predict resistance. Instead, structural variation affecting LRR and C-JID regions suggests that allele-level diversity within conserved NLR loci contributes to CR-associated variation, with implications for allele-specific marker development and durable CR deployment.

## INTRODUCTION

Clubroot is a soil-borne disease that threatens the production of the oilseed crop canola (Brassica napus L.) worldwide, especially in Canada. Yield losses in clubroot-infested canola fields can range from 30% to 100% (Botero-Ramírez et al., 2022). This disease is caused by the obligate biotrophic protist Plasmodiophora brassicae (Adhikary et al., 2025; Javed et al., 2023). Roots infected by P. brassicae become swollen and distorted into large, club-shaped galls, reducing the host’s capacity to absorb water and nutrients and leading to a substantial yield loss (Storfie et al., 2025). The life cycle of P. brassicae consists of two stages: the primary infection in both root hairs and epidermal cells, and the secondary infection in cortical cells (Liu et al., 2020). As infected roots break down, resting spores are released into the soil, where they can survive for up to 20 years, making clubroot management difficult once the field is infected (Dixon, 2009).

Compared with other management strategies, deploying resistant (R) genes through the development of clubroot-resistant (CR) cultivars remains the most effective and sustainable approach for controlling clubroot (Javed et al., 2023). Over the past two decades, extensive efforts have identified diverse sources of clubroot resistance across Brassica species, with European fodder turnips (Brassica rapa) serving as the primary donor of resistance used in commercial breeding programs (Yang et al., 2022). These efforts have led to the mapping of more than 50 quantitative trait loci (QTLs), many of which are concentrated on chromosomes A03 and A08 (Lai et al., 2025; Wei et al., 2025; Xu et al., 2025). As a result, most commercial canola cultivars now carry one or more introgressed CR genes, including both first-generation cultivars containing a single resistance gene and second-generation cultivars that pyramid multiple resistance genes. In Canada alone, more than 74 CR cultivars have been registered or commercialized in recent years (Wu et al., 2026). Despite this remarkable breeding success, the molecular basis of resistance in these cultivars remains largely unknown. Progress has been hindered because most commercially deployed CR genes are proprietary assets held by seed companies, limiting access to the germplasm and associated genotypic information (Botero-Ramirez et al., 2024). Furthermore, many CR genes provide pathotype-specific resistance, making them vulnerable to the rapid emergence of resistance-breaking P. brassicae populations (Fredua-Agyeman et al., 2021). For example, nearly 90% of isolates collected in Alberta in 2023 were able to overcome first-generation clubroot resistance (Storfie et al., 2025). As resistance-breaking populations continue to spread, identifying and characterizing new CR genes has become a priority for developing cultivars with more durable and broadly effective resistance.

Plants utilize their innate immune system to combat pathogens, and immune receptors of the nucleotide-binding leucine-rich repeat (NLR) family can recognize pathogen-derived effectors, triggering effector-triggered immunity (ETI) and thereby inhibiting pathogen spread from the site of infection (Jones et al., 2024; Ngou et al., 2022). NLR receptors play a central role in clubroot resistance, and several CR genes against P. brassicae have been cloned from different Brassica species, such as CRa, Crr1a, and Bol.TNL.2 (Hatakeyama et al., 2013; Shi et al., 2025; Ueno et al., 2012). Canonical NLRs in plants consist of three domains: an N-terminal domain, a central nucleotide-binding (NBARC) module, and a C-terminal leucine-rich repeat (LRR) domain (Barragan & Weigel, 2021). NLRs are typically classified into three main groups based on their N-terminal domains as Toll/interleukin 1 receptor (TIR) NLR (TIR-NLR), Rx-type coiled-coil NLR (CC-NLR), and resistance to powdery mildew 8 (RPW8)-type CC NLR (RPW8-NLR or CC_R_-NLR). Upon NLR activation, the N-terminal domains act as signalling domains, mediating downstream immune responses. The conserved NBARC domain serves as a switch of the NLR receptor, and the LRR domain is highly variable and functions in both direct and indirect detection of pathogen effectors, stabilizing the active and inactive states of the NLR receptor (Contreras et al., 2023; Zhang et al., 2026). Some TIR-NLRs have an additional C-terminal jelly roll/Ig-like (C-JID) domain, located after the LRR, which can strengthen LRR-effector binding (Ma et al., 2020). Additionally, some sensor NLRs with an extra non-canonical domain that does not share their evolutionary history are referred to as NLRs with integrated domains (NLR-IDs) (Marchal et al., 2022; Sarris et al., 2016). NLR receptors exhibit a high degree of structural and functional diversity because they evolve under intense pathogen-imposed selection and possess a modular architecture, which enables plants to recognize a broad range of pathogen effectors while activating distinct downstream immune responses (Chakraborty & Ghosh, 2020; Jacob et al., 2013; Monteiro & Nishimura, 2018).

Accurate NLR annotation in canola (Brassica napus) is particularly challenging owing to the complexity of its allotetraploid genome (AACC, 2n=4x=38) and high genomic redundancy (Chalhoub et al., 2014; Cheng et al., 2025). Resistance gene enrichment sequencing (RenSeq)-based enrichment can help address these limitations by providing high-depth, NLR-focused sequence data that resolves paralogous copies and identifies novel or divergent NLR genes absent from existing reference annotations (Wu et al., 2024). In addition, NLR genes also show extensive allelic and presence/absence variation in plant populations (Barragan & Weigel, 2021). By profiling NLR repertoires (NLRome) across diverse accessions or cultivars, researchers can capture variations and allelic diversity that a single reference genome would miss (Lin et al., 2023; Van de Weyer et al., 2019), which can enhance our understanding of disease resistance mechanisms and facilitate the effective utilization of high-quality R gene resources. Recently, the first comprehensive pan-NLRome comprising 23 ecotypes of B. napus was constructed, revealing variation in the distribution, expression, and evolution of NLR genes across ecotypes (Ning et al., 2024). However, there is no pan-NLRome that includes B. napus CR commercially bred cultivars, which limits our understanding of the full spectrum of resistance-associated NLR diversity, including rare or cultivar-specific genes, their evolutionary dynamics, and their contributions to durable resistance against the clubroot pathogen.

To address this knowledge gap, we combined whole-genome sequencing (WGS) and RenSeq to assemble and annotate the NLRome of five inbred homozygous (IH) B. napus lines with distinct clubroot resistance profiles. We then integrated these datasets with the universally susceptible cultivar Westar to construct a pan-NLRome that captures extensive NLR diversity across contrasting genetic backgrounds. This resource reveals patterns of NLR presence–absence and allelic variation in CR breeding lines, providing new insights into the genetic architecture of clubroot resistance and highlighting the importance of allele-level diversity for the development of more durable CR cultivars.

## RESULTS

### Distinguishable phenotypes of five CR lines to different isolates

The five homozygous CR inbred lines obtained from Nutrien Ag Solutions (Saskatoon, SK, Canada) comprise three lines carrying single-gene resistance (SGCR; IH1, IH2, and IH5) and two lines carrying multi-gene resistance (MGCR; IH3 and IH4) (Figure 1A, Table S2). Their resistance profile was evaluated against four P. brassicae pathotypes (3H, 3D, 3A, and 5X), which were defined using the Canadian Clubroot Differential (CCD) set and are among the predominant pathotypes in Western Canada (Hollman et al., 2023; Storfie et al., 2025). According to our industrial partners, IH1 and IH2 carry a CR source that confers resistance to pathotype 3H. IH1 carries a CR gene derived from the European winter rapeseed cultivar “Mendel”, whereas IH2 originates from Chinese cabbage (Brassica rapa). Previous genetic mapping indicates that these two lines share a common CR locus on chromosome A03. In contrast, IH3, IH4, and IH5 are considered second-generation CR lines harboring either novel or stacked resistance traits, reportedly derived from rutabaga (B. napus). The IH3-IH5 lines were developed by introgressing a novel CR source from rutabaga, and a series of crosses and backcrosses with different germplasms. IH3 and IH4 possess a CR loci on both A03 and A08 and exhibit resistance to all four pathotypes tested, whereas IH5 contains a locus on A08 only and shows a narrower resistance profile, being resistant to 3D and 5X but susceptible to 3A and 3H (Figure 1A, Table S2).

**FIGURE 1.**
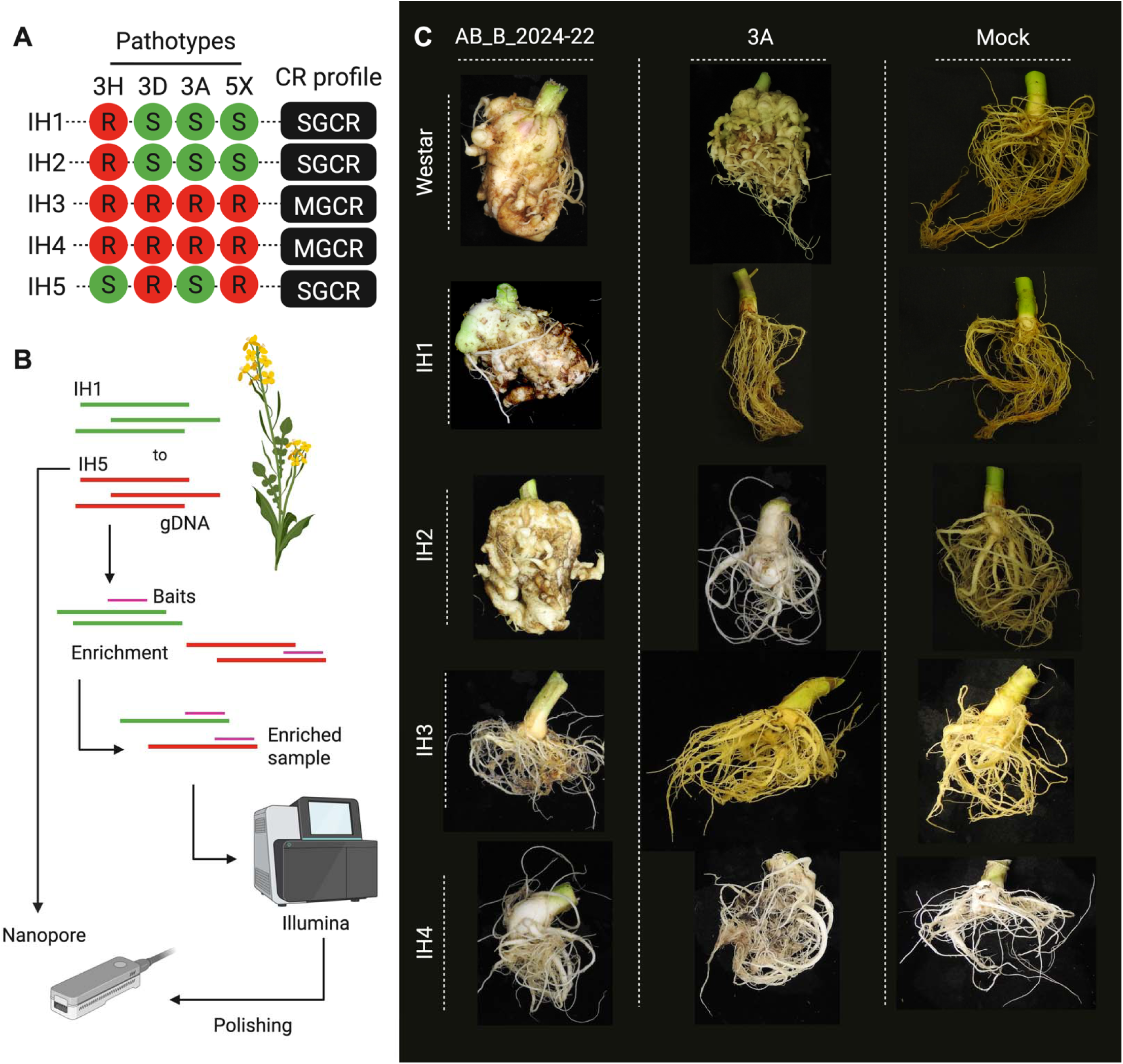
Identification of clubroot resistance profiles and genome assembly in Brassica napus. **(A)** Clubroot resistance (CR) profiles of five inbred homozygous canola lines (IH1 to IH5) against four P. brassicae pathotypes (3H, 3D, 3A, and 5X). Red circles indicate CR, and green circles indicate clubroot susceptibility (S). Based on their resistance profile and breeding history, lines are classified as single gene clubroot resistance (SGCR) or multi-gene clubroot resistance (MGCR). (**B**) Schematic overview of the RenSeq and whole-genome sequencing workflow. Genomic DNA from the CR lines was subjected to bait capture targeting NLR sequences and to Illumina sequencing. Meanwhile, gDNA was also used for Oxford Nanopore sequencing. The raw RenSeq reads were used to polish the reads during genome assembly. (**C**) Representative root phenotypes of Westar (susceptible control) and IH1 to IH4 lines following inoculation with P. brassicae pathotype AB_B_2024-22 (highly virulent) and 3H (less virulent), alongside mock-inoculated controls. Severe gall formation is observed in susceptible hosts, whereas CR lines display reduced or no galling and maintain relatively normal root architecture depending on the isolated used to challenge the canola line. Differences in symptom severity across genotypes and pathotypes highlight distinct resistance profile among the tested lines.

To validate these resistance profiles, we performed root phenotyping assays with two additional P. brassicae isolates under controlled conditions. When challenged with pathotype 3H and the isolate AB_B_2024-22, which exhibits virulence comparable to pathotype 5X, and evaluated at 42 days post-inoculation (dpi), the SGCR lines IH1 and IH2 exhibited strong resistance to pathotype 3H but were highly susceptible to AB_B_2024-22, with severe gall formation similar to the susceptible control Westar (Figure 1C). The MGCR lines IH3 and IH4 maintained robust resistance to both, with minimal or no gall development and largely intact root architecture (Figure 1C). Additionally, the SGCR line, IH5, shows partial resistance to the 3H pathotype and resistance to the AB_B_2024-22 isolate (Figure S1). These results highlight that stacked genetic clubroot resistance exhibits broader effectiveness against P. brassicae infection compared with single-gene resistance.

### WGS and RenSeq to reveal the NLRome of CR canola lines

NLR immune receptors are well established as central components of disease resistance, and at least two cloned CR genes have been identified as members of the NLR family (Wu et al., 2026). To further dissect the genetic basis of CR in this germplasm, we combined WGS and RenSeq (Figure 1B). We first constructed the pipeline to de novo assemble the genomes of these five lines and annotate the NLR receptors (Figure S2). Through WGS, we achieved ∼10× coverage, while RenSeq provided >250× coverage in NLR-enriched regions.

Across all CR lines, a comparable number of NLR genes were identified, ranging from 504 in IH5 to 517 in IH2 (Figure 2A). These NLRs were classified into full NLRs (TIR-NLR, CC-NLR, and RPW8-NLR) and their corresponding partial domain architectures (TIR-NBARC, CC-NBARC, RPW8-NBARC, NBARC-LRR, and NBARC) (Table S3). Additionally, the number of NLRs in each class across these five CR lines was comparable (Table S3). Among the full NLRs, TIR-NLRs are the predominant class across all lines, which is consistent with previous observations in Brassica species, followed by CC-NLRs and RPW8-NLRs (Ning et al., 2024) (Figure 2A, Table S3). Despite the overall similarity in total NLR counts, differences were observed in the proportion of intact versus partial NLRs among lines (Figure 2B, Table S4). Specifically, IH3 harbours the highest number of partial NLRs (251), whereas IH5 contains the lowest (230). The number of full-length NLRs is relatively stable across genotypes, ranging from 264 to 275 across lines (Figure 2B).

**FIGURE 2.**
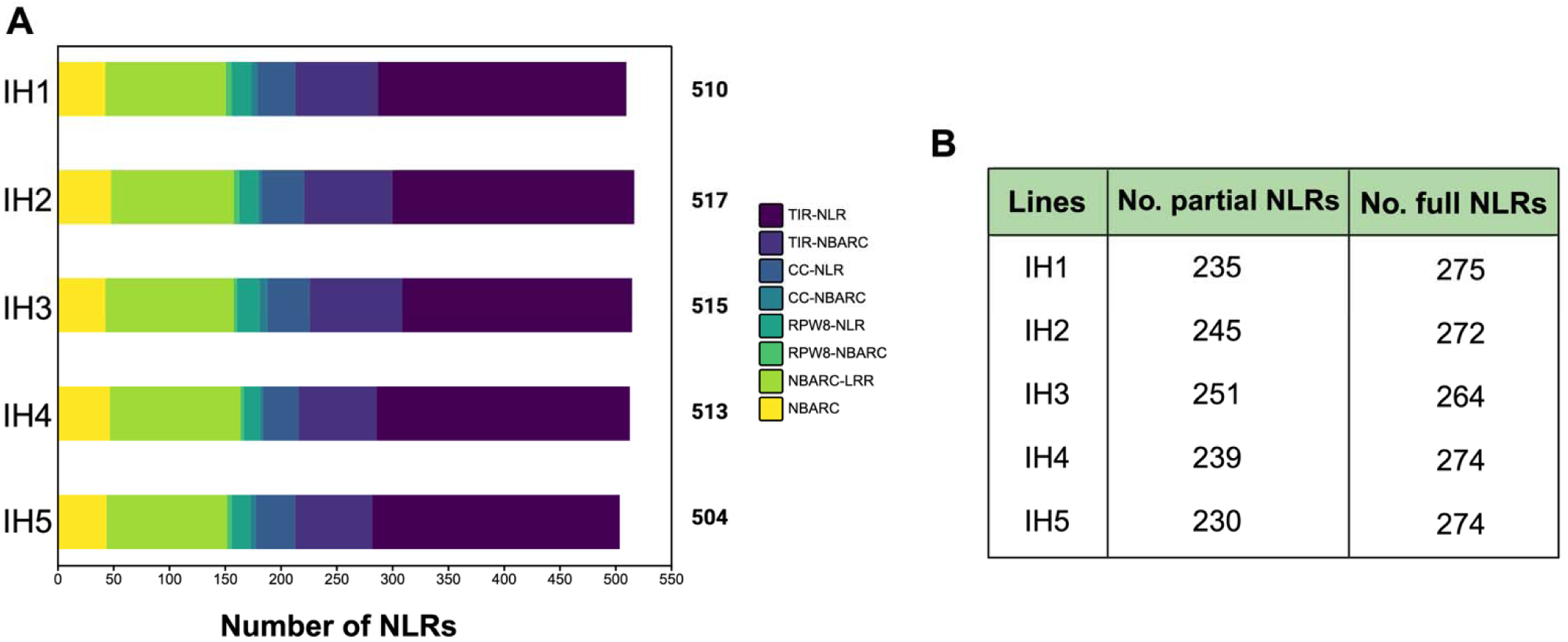
NLR gene repertoire across five inbred homozygous canola lines. **(A)** Stacked horizontal bar chart showing the total number of NLR genes identified in each inbred line (IH1–IH5), classified by domain architecture: TIR-NLR, TIR-NBARC, CC-NLR, CC-NBARC, RPW8-NLR, RPW8-NBARC, NBARC-LRR, and NBARC, with TIR-NLR representing the predominant classes. Total NLR counts per line are indicated to the right of each bar. (**B**) Table summarizing the number of partial and full-length NLR genes identified in each line. Across all five lines, NLR complements were broadly conserved, ranging from 504 (IH5) to 517 (IH2).

The NBARC domain is the most conserved part of an NLR immune receptor and is useful for determining evolutionary relationships among NLRs through global alignments (Kourelis et al., 2021). To study the evolutionary relationships among the NLR genes of the CR lines, we extracted the NBARC domain (PF00931) from each NLR protein to construct a maximum-likelihood phylogenetic tree. Phylogenetic analysis revealed a structured clustering of NLRs into distinct clades corresponding to major subclasses (Figure 3A). Clear separation of canonical NLR types, including CC-NLR and TIR-NLR groups, was observed, indicating conserved evolutionary relationships across all five genotypes (Figure 3A).

**FIGURE 3.**
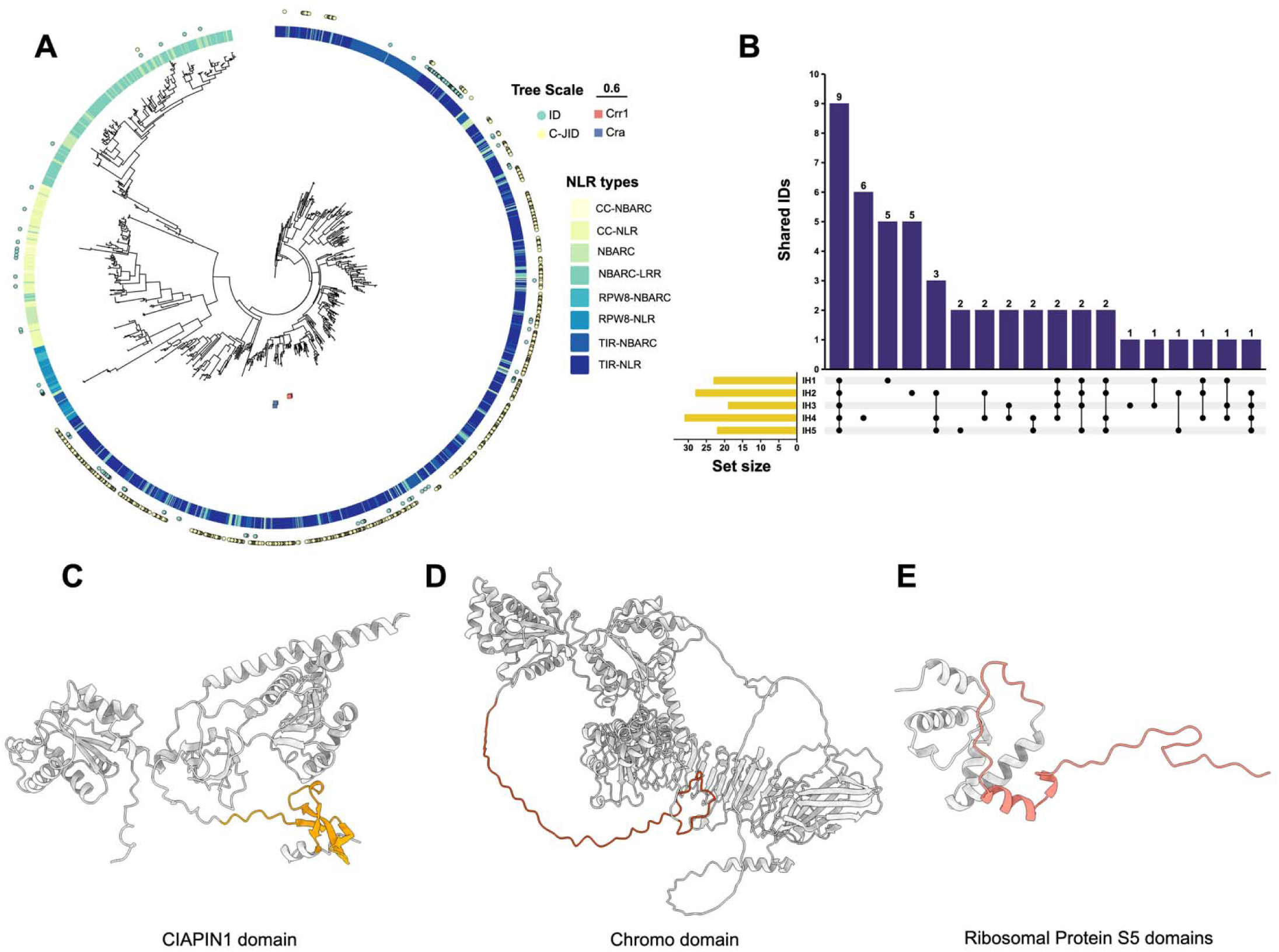
NLRome architecture and Integrated domain (ID) diversity across five clubroot-resistant canola lines. **(A)** The NBARC domain sequences were extracted and used to construct a maximum-likelihood phylogenetic tree of NLR proteins identified in five CR lines (IH1 to IH5). The outer rings indicate NLR subclasses, annotated by domain architecture (e.g., CC-NLR, TIR-NLR, and other subclasses). IDs and C-JID domains fused to NLR proteins are indicated by green and yellow circles, respectively. The CRa and Crr1a homologs were indicated by blue and pink squares, respectively. Tree scale is 0.6. (**B**) UpSet plot showing the distribution and intersections of NLR-IDs across the five lines. The bar chart (top) shows the number of shared or unique NLR-IDs across genotype combinations, and the connected dots (bottom) indicate the specific sets of lines contributing to each intersection. The horizontal bars (left) show the total number of NLR-IDs identified by each line. (**C-E**) The protein structures with unique NLR-IDs include the CIAPIN domain, Chromo Domain, and Ribosomal Protein S5 domain. These structures were generated by AlphaFold3 (Abramson et al., 2024).

The presence of NLR-IDs indicates a highly evolved and versatile immune surveillance system in the plant host. We next examined the distribution of IDs within NLRs across the lines (Table S5). Nine IDs were shared among all lines (Figure 3A, B), including well-characterized domains such as heavy-metal-associated (HMA), zinc finger, LIM, and B3 DNA-binding domains, highlighting the conservation of NLR-IDs. Moreover, the IH4 line contained the highest number of unique IDs, several of which have not been previously reported, including chromo, cytokine-induced anti-apoptosis inhibitor 1 (CIAPIN), and ribosomal protein S5 domains (Figure 3C-E). Additionally, the C-JID domain was specifically fused to the C-terminus of TIR-NLRs, a configuration known to enhance LRR-mediated effector recognition (Ma et al., 2020). C-JID-containing NLRs accounted for more than 30% of the total NLR repertoire in each CR line (Figure 3A, Table S4).

### NLRs among CR and susceptible canola lines remain largely conserved

To identify resistance-associated NLR genes and capture the broader diversity of the canola NLRome, we constructed a pan-NLRome integrating the CR lines generated in this study with the previously reported NLRome of the susceptible cultivar Westar (Wu et al., 2024). This comparative framework enabled the identification of candidate CR-associated NLRs by distinguishing resistance-specific variation from the background NLR diversity present across B. napus genomes. Firstly, through the pairwise reciprocal best hits (RBH) across these six lines, we found that the average number of RBH pairs between 5 CR lines is 362 (range 340-400). In contrast the average number of RBH pairs per CR line to Westar is 302 (range 285-321), indicating that the five CR lines are noticeably closer to each other than to susceptible Westar. Moreover, the IH1 and IH2 have the highest NLRome similarity, which contains 400 RBH pairs (Figure 4A, Table S6).

**FIGURE 4.**
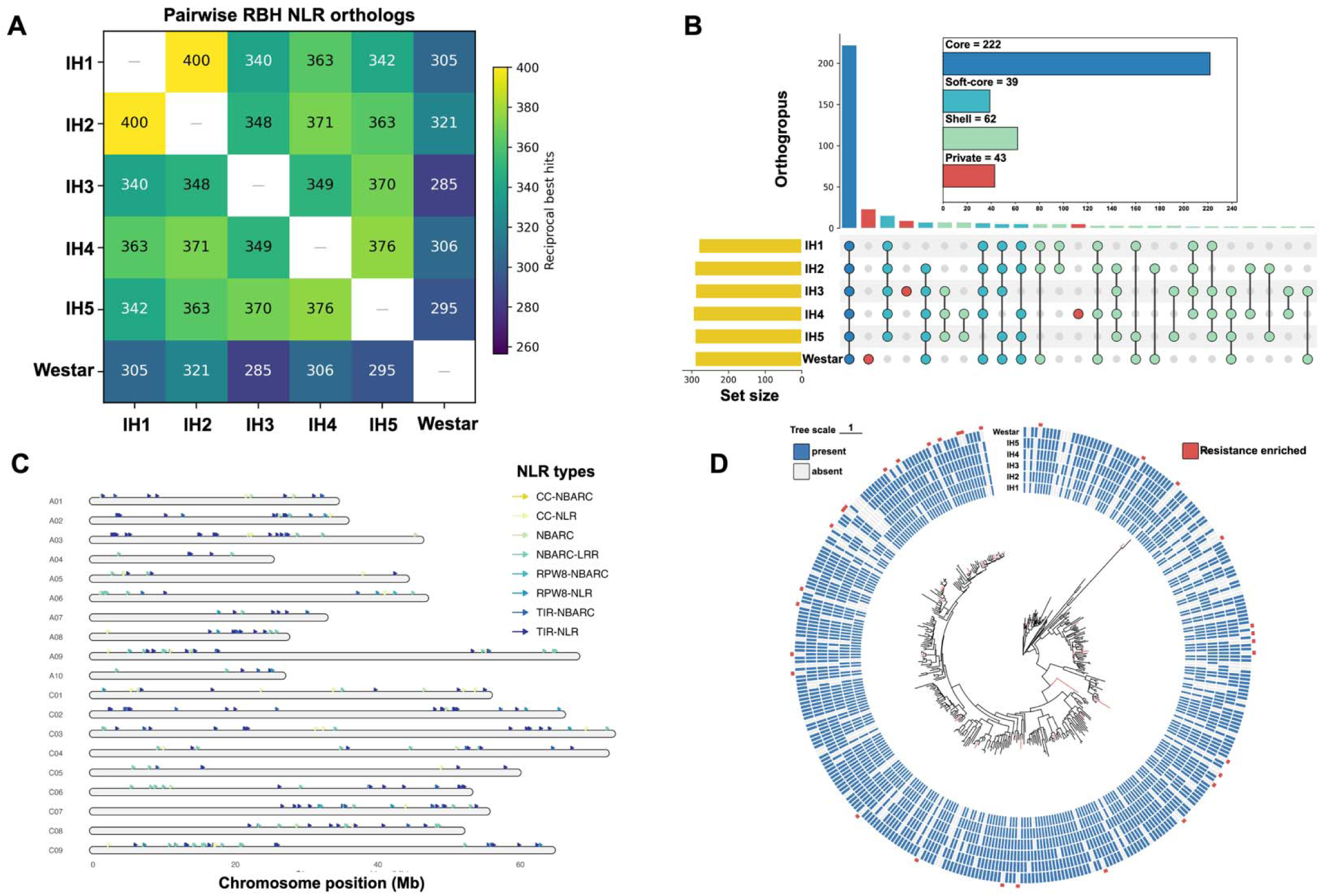
Pan-NLRome architecture, distribution, and phylogeny across five CR canola lines and Westar. **(A)** Pairwise reciprocal best-hit (RBH) NLR orthologs. Heatmap of the number of one-to-one NLR pairs identified by MMseq2 easy-RBH between every pair of lines. (**B**) UpSet plot showing the distribution of NLR OGs across the six resistant lines. The bar chart (top) indicates the number of Ogs shared among different combinations of lines, while the matrix below denotes the specific intersections. The horizontal bar plot (left) represents the total number of NLR OGs detected in each line. Classification of NLR OGs into core, shell, and private components based on their presence across resistant lines. Core OGs are present in all six lines (6/6, n = 222), soft-core OGs are present in five lines (5/6, n = 39), shell OGs are present in two to four lines (2-4/6, n = 62), and private OGs are unique to a single line (1/6, n = 43). (**C**) Genome-wide distribution of the 464 Westar genes assigned to core OGs across 19 canola chromosomes. Different NLR classes are indicated by distinct colors. Chromosome positions are shown along the x-axis (Mb). (**D**) Phylogenetic tree of NBARC domains representing the pan-NLRome, combined with NLR presence/absence matrix across cultivars. The maximum likelihood tree was constructed using representative NBARC sequences from each OG. The outer heatmap indicates the presence (blue) or absence (light gray) of each OG in individual canola lines. Red marks highlight resistance-enriched OGs (present in ≥ 3 R-lines and absent in Westar). Tree scale is 1.

Next, we clustered the NLR genes from six lines into orthogroups (OGs) based on sequence similarity and orthology inference to build the pan-NLRome. The species tree shows Westar is a clean outgroup with 100% bootstrap support, and the five resistant lines form two well-supported subclades (IH1/IH2 and IH3/IH4/IH5) (Figure S3). In total, 366 NLR OGs were identified (Table S7, Figure 4B). These OGs were further classified into four categories based on their distribution across accessions as (i) core, (ii) soft-core, (iii) shell, and (iv) private (Figure 4B; Tables S8 and S9). The core NLRome comprised 222 OGs (60.7%) and represented 2,660 genes present in all six germplasm. The soft-core NLRome contains 39 OGs and 230 genes across five accessions. The shell NLRome included 62 OGs containing 177 genes shared by two to four accessions, while the remaining 43 OGs (11.7%) were classified as private, comprising 47 genes detected in only a single line (Figure 4B; Tables S8 and S9).

To examine the genomic organization of NLRs based on Westar reference genome, we mapped 464 core NLR genes across all B. napus (Westar) 19 chromosomes (Figure 4C). NLR genes displayed a highly uneven distribution, forming distinct OGs in specific chromosomal regions. Several chromosomes, including A03 and A08, contained high densities of NLRs, whereas others harbored relatively few genes, indicating a non-random genomic arrangement (Figure 4C; Table S10). This pattern suggests that local duplication and expansion events show dynamic evolution of NLR gene families, which have contributed to the accumulation of NLR genes in particular genomic regions.

To further investigate how conserved or variable the NLRs are between CR and the susceptible canola lines, we constructed a maximum-likelihood phylogenetic tree using the NB-ARC domains of all 366 NLR OGs and overlaid OGs presence-absence patterns across the six germplasm lines (IH1 to IH5 and Westar) (Figure 4D). The majority of NLRs were conserved across all lines and identified as the core set, supporting the predominance of the core NLRome. However, a subset of OGs showed lineage-specific absence, indicating diversification among the resistant lines. There is a total of 75 OGs present only in the five CR lines but absent in Westar (Figure S4A, Table S11). Based on the different types of their CR profiles, we found that 9 OGs are present in the first-generation SGCR group. There are 16 OGs associated with the second-generation MGCR, with the larger NLR diversity. However, the OGs only present in IH5 are small but distinct second-generation SGCR-specific signals (Table S11). More importantly, 31 OGs contained NLRs enriched in at least 3 resistant lines but absent in Westar, suggesting that these NLRs are potential candidates for clubroot resistance (Figure 4D, Figure S4A, Table S12).

### Allelic variation of clubroot resistance genes CRa and Crr1a among CR lines

The pan-NLRome analysis indicated high conservation of NLR across CR and susceptible lines. To further interpret the pan-NLRome, we first assessed the representation of two known CR genes (CRa and Crr1a). The CRa locus, originally identified on chromosome A03 in B. rapa, encodes a well-characterized TIR-NLR protein conferring CR (Ueno et al., 2012). Based on the source of introgression, IH1-IH4 were expected to carry CRa, whereas IH5 was not. To verify this, the reference CRa protein sequence (GenBank accession: BAN04701.1) was used to query the five NLRomes. Homologs of CRa were identified in IH1 to IH4 (IH1_contig_4441_000164.1, IH2_contig_1104_000037.1, IH3_contig_4607_000008.1, and IH4_contig_1572_000024.1) (Figure 5A-D, Figure S5), with high sequence RenSeq-coverage read support across the locus. Surprisingly, a CRa-like sequence was also detected in IH5 (IH5_contig_3416_000002.1) (Figure 5E; Figure S5), suggesting a broader distribution of this gene than previously expected. Synteny analysis between the CR loci CRA3.7 indicates broad conservation of the genomic neighbourhood surrounding CRA3.7.1/CRa between ECD04 and the five lines. Multiple NLR genes flanking CRa were connected by conserved syntenic relationships, while IH1 and IH2 showed relatively similar local organization to the ECD04 reference, whereas the regions in IH3–IH5 were more divergent (Figure 5F). Multi-sequence alignment revealed a 135-bp deletion within the LRR-encoding region in IH3 and IH4, resulting in a 45-amino-acid truncation (Figure 5G). We used AlphaFold 3 to model the structures of the two CRa alleles. The structural comparison showed that the deletion is located in an LRR domain (Figure 5H). These results indicated that CRa homologs are broadly present among the resistant lines but display distinct LRR allelic variation.

**FIGURE 5.**
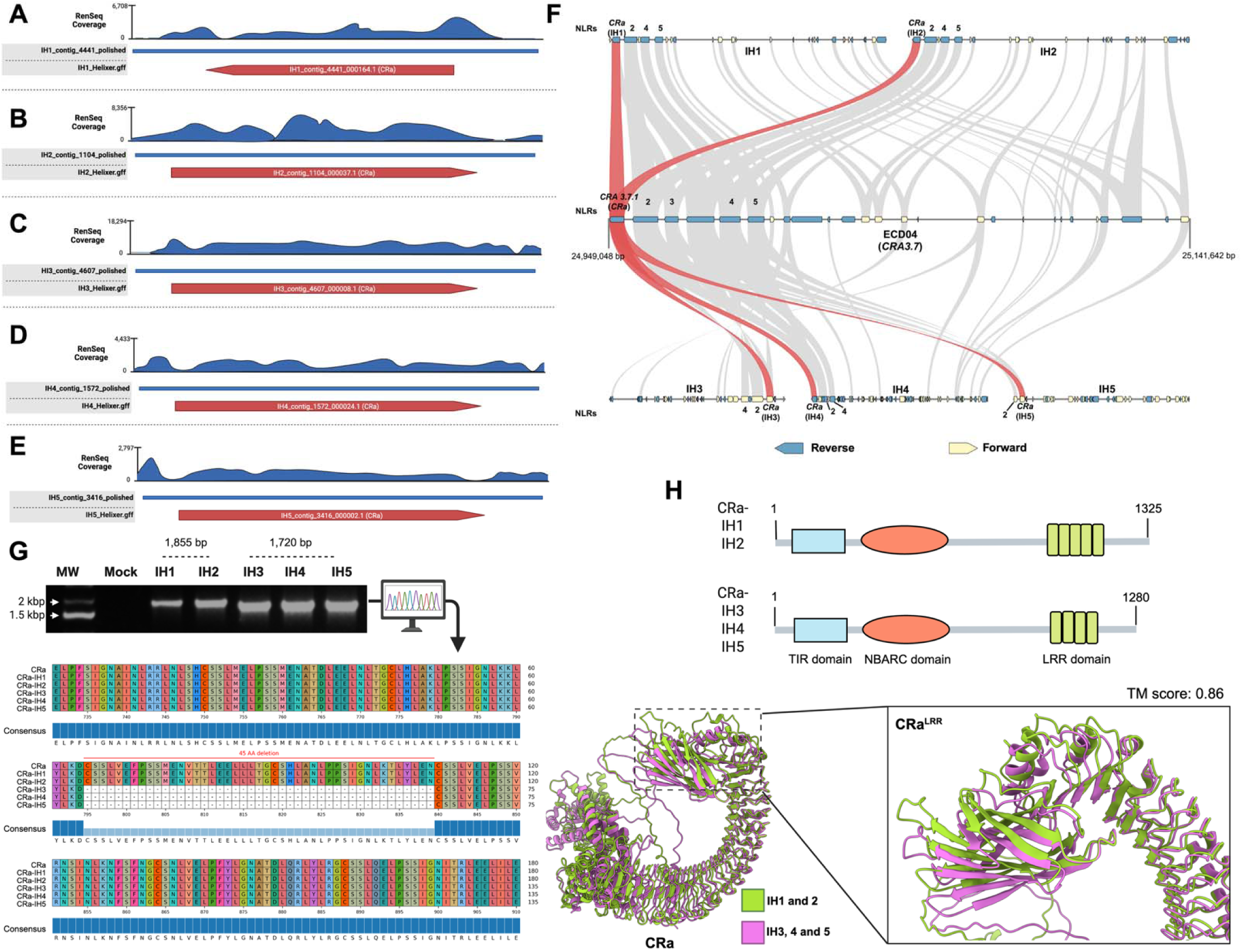
RenSeq-based validation and structural variation of CRa homologs across five CR lines. **(A-E)** RenSeq read coverage mapped to the assembled genomes of five resistant lines (IH1 to IH5) at the CRa locus. Blue tracks represent sequencing coverage across each contig, and red arrows indicate predicted CRa gene models. Enrichment and full-length coverage across all lines support the successful assembly and presence of CRa-like sequences. The read depth for each CRa homolog gene was visualized using Geneious Prime. (**F**) Synteny between the CRa candidate regions. Global synteny of the CR loci on A03 (CRA 3.7) of ECD04 and the candidate regions of each IH line. The number above the arrow boxes emphasizes the NLR genes in the region. Yellow and blue arrow boxes represent the orientation of the coding sequences (CDS) of all genes. Grey shadow areas represent conserved syntenic blocks, and the red ribbon shows the position of the CRa homologs. (**G**) PCR amplification of the LRR region CRa from IH1 to IH5 confirms its presence in all lines, with two distinct amplicon sizes (1,855 bp in IH1 and IH2, and 1,720 bp in IH3 to IH5), suggesting structural variation. Multiple sequence alignment of the encoded proteins highlights overall conservation, except for a 45-amino-acid deletion in the LRR domain. (**H**) Schematic representation of the domain architecture and structures of CRa proteins identified in the five lines. CRa homologs from IH1 and IH2 encode a longer protein of 1,325 amino acids, whereas CRa homologs from IH3, IH4, and IH5 encode a shorter protein of 1,280 amino acids. Predicted structural models of the alignment of CRa alleles show that the deletion is within the LRR domain. Green indicates CRa alleles from IH1 and IH2, and magenta indicates CRa alleles from IH3, IH4, and IH5.

Next, we examined another well-characterized clubroot resistance gene, Crr1a, which also encodes a TIR-NLR and is localized on chromosome A08 (Hatakeyama et al., 2013; Hatakeyama et al., 2022). The Crr1a gene is expected to be present in lines IH3-IH5. Based on the results of reference Crr1a (GenBank accession: BAM77406.1) query to five NLRomes. Homologs of the Crr1a gene were found in all five CR canola lines (IH1_contig_1575_000235.1, IH2_contig_4267_000144.1, IH3_contig_3767_000202.1, IH4_contig_1467_000016.1, and IH5_contig_3607_000160.1) (Figure 6A-E). Synteny analysis of the CRA8.2 region revealed conservation of the genomic interval containing CRA8.2.4/Crr1a between ECD04 and the five lines. While differences in gene order, spacing, and local gene composition were evident among the accessions, indicating structural diversification of the Crr1a-containing region (Figure 6F). Moreover, we found that the sequence in the IH1 and IH2 lines lacks 230 amino acids in the C-terminal region compared to IH3, IH4, and IH5 (Figure 6G-H). Domain architecture analysis further showed that this variation affects the LRR region, and additionally, IH1 and IH2 lack the C-terminal C-JID domain present in IH3 to IH5 (Figure 6G). Structural comparison of these Crr1a homologs further revealed substantial differences in the LRR/C-JID region between the two allele groups (Figure 6H), which might influence recognition specificity or downstream immune function and could contribute to the different clubroot resistance profiles observed among these lines. Both CRa and Crr1a were assigned to the core NLR OGs in the pan-NLRome, which are OG0000051 and OG0000014, respectively. Their widespread conservation across both resistant and susceptible backgrounds indicates that resistance is unlikely to be explained by gene presence/absence alone. Instead, these results suggest that allelic variation, particularly within the LRR domain, may underlie functional divergence through effector recognition.

**FIGURE 6.**
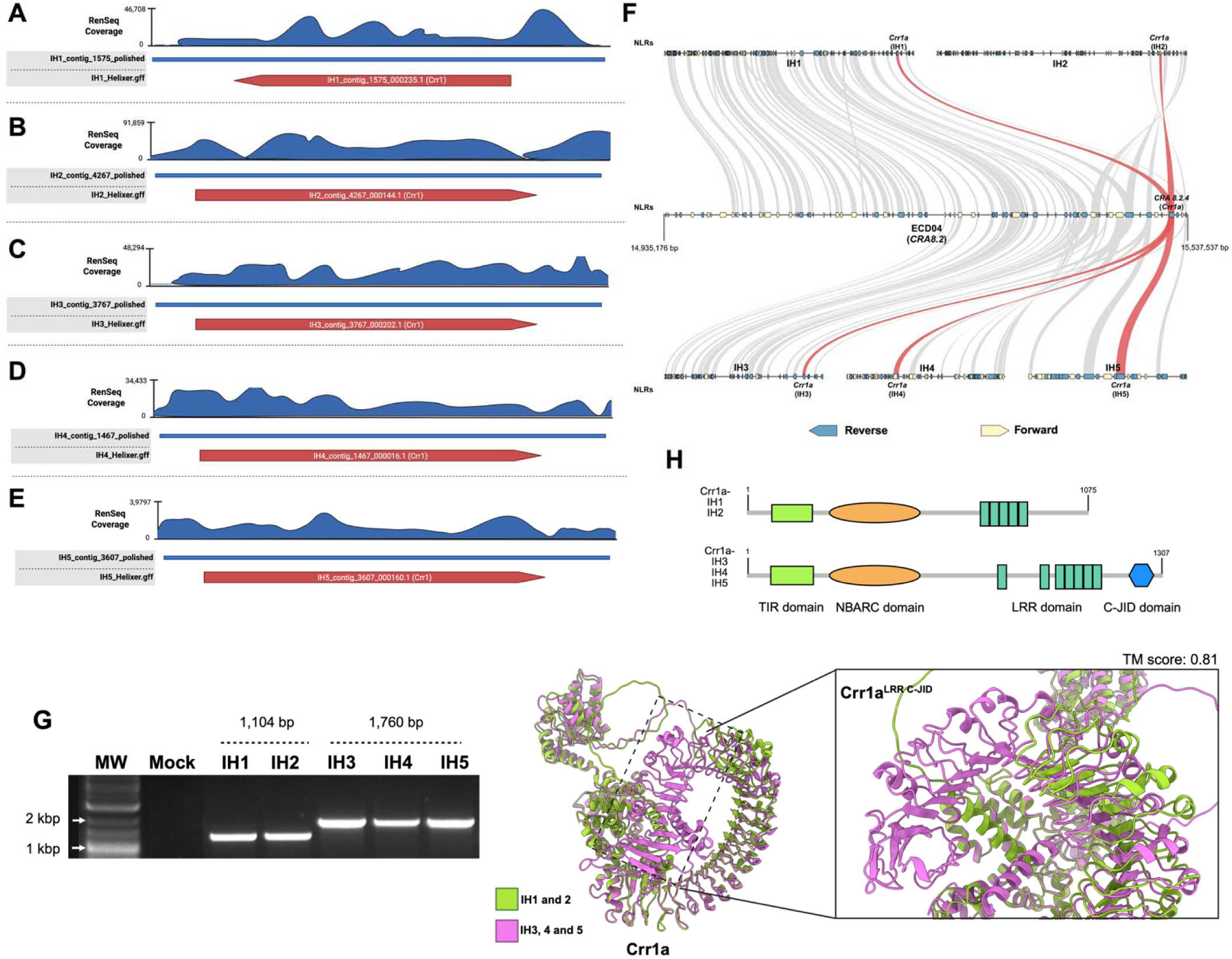
RenSeq-based identification and structural variation of Crr1a homologs across five clubroot-resistant lines. **(A-E)** RenSeq read coverage mapped to the assembled genomes of five resistant lines (IH1 to IH5) at the Crr1a locus. Blue tracks represent sequencing depth across each contig, and red arrows indicate predicted Crr1a gene models. Enrichment and continuous coverage across all lines support the assembly and presence of Crr1a sequences. (**F**) Synteny between the CRa candidate regions. Global synteny of the CR loci on A08 (CRA 8.2) of ECD04 and the candidate regions of each IH line. Yellow and blue arrow boxes represent the orientation of the coding sequences (CDS) of all genes. Grey shadow areas represent conserved syntenic blocks, and the red ribbon shows the position of the Crr1a homologs. (**G**) PCR amplification of Crr1a confirms its presence in all five resistant lines, with two distinct amplicon sizes (1,104 bp in IH1 and IH2 and 1,760 bp in IH3 to IH5), indicating structural differences among the alleles. These variations in domain composition, particularly in the LRR and C-terminal regions, may contribute to functional divergence in clubroot resistance. (**H**) Schematic representation of the domain architecture and structures of Crr1a proteins identified in the five lines. All proteins contain conserved TIR and NBARC domains; however, structural variation is observed in the C-terminal region. IH1 and IH2 encode canonical Crr1a proteins with only an LRR domain, whereas IH3, IH4, and IH5 exhibit a modified structure with alterations in the LRR region and the presence of an additional C-JID domain. Predicted structural models of the alignment of Crr1a alleles show that the deletion is within the C-terminal. Green indicates Crr1a alleles from IH1 and IH2, and magenta indicates Crr1a alleles from IH3, IH4, and IH5.

## DISCUSSION

Brassica napus is a relatively young allotetraploid species that originated approximately 7,500 years ago through hybridization between B. rapa (AA, 2n = 20) and B. oleracea (CC, 2n = 18) (Chalhoub et al., 2014). Due to the recent origin and narrow genetic diversity of cultivated B. napus, ancestral and related Brassica germplasm has played a critical role as a reservoir of R genes against multiple pathogens, including Plasmodiophora brassicae, the causal agent of clubroot disease. In this study, we analyzed five commercial CR canola breeding lines carrying resistance derived from different genetic backgrounds, including rapeseed cv. Mendel (IH1), Chinese cabbage (IH2), and rutabaga cv. Brookfield (IH3–IH5) (Table S2). Interestingly, many of these resistance sources ultimately trace back to European fodder turnip accessions, highlighting the importance of related Brassica germplasm in expanding the immune repertoire available for canola improvement (Fredua-Agyeman & Rahman, 2016; Hasan & Rahman, 2016; Hatakeyama et al., 2022; Yang et al., 2022). Previous studies suggest that some CR genes originated from a common ancestor before the whole-genome triplication event, indicating that current resistance diversity may reflect the retention and redistribution of ancient immune receptors rather than recent gene emergence (Yang et al., 2022). Consistent with this hypothesis, the diverse resistance sources observed across the five breeding lines suggest that clubroot resistance in Brassica crops has largely been shaped by the selection and deployment of pre-existing NLR diversity rather than the evolution of lineage-specific R genes (Durand et al., 2020; Gossen et al.; Lai et al., 2025; Lv et al., 2020).

Most CR canola cultivars exhibit pathotype-specific resistance, making them vulnerable to resistance breakdown under field conditions, particularly when protection relies on a single major R gene (Fredua-Agyeman et al., 2018). This limitation became evident following the widespread deployment of first-generation CR cultivars, when resistance-breaking pathotypes such as 3A and 5X emerged and became prevalent across Canadian canola-growing regions (Hollman et al., 2023; Storfie et al., 2025). Our phenotyping results showed that the first-generation CR lines IH1 and IH2 were resistant to pathotype 3H but susceptible to the more virulent isolate AB_B_2024-22 (Figure 1C). To improve resistance durability, second-generation CR cultivars carrying additional sources of resistance were introduced in Canada in 2019. In agreement with their broader resistance spectrum, IH3 and IH4 showed strong resistance to both 3H and AB_B_2024-22, whereas IH5 was resistant to AB_B_2024-22 but displayed only partial resistance to 3H (Figure 1C, Figure S1). These contrasting phenotypic responses suggest that distinct genetic mechanisms underlie resistance in first- and second-generation CR lines and highlight the importance of matching cultivar deployment with local P. brassicae pathotype composition for effective and durable clubroot management.

The NLR family is one of the most diverse gene families in plants, with NLR repertoires varying extensively across species and genome architectures (Barragan & Weigel, 2021; Wu et al., 2026). Given the complexity and repetitive nature of NLR loci, RenSeq has become a powerful approach to improve NLR annotation and construct comprehensive NLRomes and pan-NLRomes across diverse plant species (Andolfo et al., 2014; Huang et al., 2022; Jupe et al., 2013; Parada-Rojas et al., 2025; Van de Weyer et al., 2019; Vendelbo et al., 2022). By integrating WGS and RenSeq, our analysis revealed that the overall NLR repertoire is highly conserved across the five CR canola breeding lines, suggesting that differences in total NLR abundance alone are unlikely to explain variation in clubroot resistance. Instead, differences in NLR composition, including the proportion of partial NLRs, suggest ongoing receptor turnover and structural diversification that may contribute to genotype-specific immune responses (Jacob et al., 2013). Although partial NLRs may lack canonical domains, such as N-terminal signalling domains or LRR regions, growing evidence indicates that truncated and non-canonical NLR architectures can retain key immune functions (Hu et al., 2025; Liu et al., 2017; Roth et al., 2017; Zhang et al., 2025). Consistent with previous reports in Brassica species, TIR-NLRs represented the predominant NLR class across all CR lines, contrasting with the NLR architecture typically observed in monocots (Chakraborty & Ghosh, 2020; Guo et al., 2025). The phylogenetic separation of major NLR subclasses and expansion of TIR-NLR diversity further support their central role in mediating immune responses against P. brassicae (Ning et al., 2024) (Figure 3A). Beyond canonical NLR variation, the widespread presence of IDs, including both conserved IDs shared across all lines and genotype-specific IDs such as those newly identified in IH4, highlights the contribution of domain architecture diversification to expanding effector recognition potential (Figure 3C–E, Table S5). From an evolutionary perspective, the high prevalence of C-JID-containing NLRs (∼30%), a domain fusion proposed to have emerged after the divergence of dicot lineages, suggests that these integrated domains may contribute to fine-tuning immune specificity in Brassica species (Maruta et al., 2026; Zeng et al., 2026).

The pan-NLRome analysis integrating resistant and susceptible B. napus lines revealed a largely conserved NLR repertoire, with most NLRs shared across genotypes regardless of their resistance phenotype (Figure 4B, Table S7–9) (Ning et al., 2024). This extensive core NLRome suggests that a substantial proportion of immune receptors is maintained across accessions, likely reflecting conserved components of the basal immune system (Contreras et al., 2023; Grund et al., 2019). Despite this overall conservation, the uneven chromosomal distribution and clustering of NLR loci indicate that local duplication and expansion events continue to shape NLR organization, consistent with patterns observed across other plant genomes (Figure 4C) (Silva-Arias et al., 2025; Winters et al., 2025). Importantly, diversity within the shell and private NLR fractions, together with the identification of OGs enriched in resistant lines but absent from the susceptible cultivar Westar, suggests that a smaller subset of variable NLRs may underlie genotype-specific clubroot resistance (Figure 4D, Figure S4, Table S11–12). Although these resistance-associated NLRs represent strong candidates for CR genes, further functional validation will be required to determine their specific contribution to P. brassicae resistance, or if they are implicated in other biotic or abiotic responses.

The two TIR-NLR clubroot resistance genes CRa and Crr1a have been functionally characterized, with CRa acting as a dominant resistance gene and Crr1a displaying incomplete dominance (Hatakeyama et al., 2022). Our analysis of these genes across resistant and susceptible B. napus genotypes provides new insights into the genetic basis of clubroot resistance and highlights the importance of allele-level variation. Although CRa and Crr1a were identified within the core NLR OGs and conserved across all genotypes, including the susceptible cultivar Westar, their presence alone was not predictive of resistance (Table S7). Instead, structural variation among CRa- and Crr1a-like alleles suggests that differences in domain architecture, particularly in the LRR and C-JID regions, may influence receptor function (Cesari et al., 2022). For CRa, IH3 and IH4 carried alleles with a 45-amino-acid LRR truncation and displayed the broadest resistance spectrum among the tested lines, whereas IH1 and IH2 carried full-length CRa-like alleles but were primarily resistant to pathotype 3H (Figure 5G-H). This pattern may indicate that structural changes within the LRR region modify recognition specificity; however, the higher CR performance in IH3 and IH4 may also result from additional resistance loci, including the stacked A08 region, rather than CRa variation alone (Hatakeyama et al., 2022; Javed et al., 2023; Yang et al., 2025). Similarly, Crr1a-like alleles in IH1 and IH2, which displayed the narrowest resistance profile in our panel, contained a 230-amino-acid C-terminal truncation affecting the LRR region and C-JID domain, potentially altering effector recognition or receptor activation (Figure 6G-H) (Ma et al., 2020). Previous comparisons of CRa (CRA3.7.1) and Crr1a (CRA8.2.4) homologs between resistant and susceptible lines also suggested that transposable element insertions can disrupt CR gene function, further supporting the idea that structural integrity, rather than gene presence alone, determines functional resistance (Yang et al., 2022). In addition, we found a deletion in the C-terminal region of the Crr1a allele in Westar compared to the IH3-IH5 alleles. However, further functional validation of this allelic diversity should be carried out.

To date, the mechanisms underlying these two NLR-mediated resistance responses remain poorly understood. Recent work by Piao et al. (2023) implicates calcineurin B-like 1.2 (CBL1.2), a Ca²⁺ sensor, in CRa-mediated resistance in B. rapa. In this model, CRa activation triggers cytosolic Ca²⁺ influx, which is sensed by CBL1.2 at the plasma membrane to relay downstream immune signalling. As with other TIR-NLRs, this response is likely dependent on the EDS1 pathway and requires a helper RPW8-NLR to activate defence responses (Contreras et al., 2023). However, the avirulence effectors of P. brassicae recognized by CRa and Crr1a, as well as the downstream signalling events that ultimately determine resistance, remain unknown. These unresolved mechanisms further emphasize the functional validation of the allele, and determining downstream immune signalling will be essential for understanding CR specificity.

## CONCLUSION

In this study, we combined WGS and RenSeq to characterize the NLRomes of five commercial CR canola inbred lines with contrasting resistance profiles and integrated them with the susceptible cultivar Westar to build a comparative canola pan-NLRome. Although the overall NLR repertoire was highly conserved across genotypes, we identified resistant-line-enriched NLR OGs, newly detected integrated domains, and substantial allele-level variation in known CR genes. The widespread detection of CRa and Crr1a homologs across resistant and susceptible backgrounds shows that the presence of known CR gene homologs alone is insufficient to predict resistance. Instead, structural variation in key domains, such as the LRR and C-JID regions, may contribute to functional divergence among alleles. These findings highlight allele-level characterization as a critical step for understanding clubroot resistance and for developing more reliable markers to support durable, pathotype-resilient CR breeding in canola.

## MATERIALS AND METHODS

### Plant materials and growth conditions

Canola (Brassica napus L.) seeds of five inbred homozygous lines with different CR profiles (IH1, IH2, IH3, IH4, and IH5) were provided by Nutrien Ag Solutions (Saskatoon, SK, Canada). The doubled haploid Westar that carries no CR gene was used as a susceptible control. Seeds were germinated in vermiculite for 5 days, and seedlings were then transplanted into soil-filled pots in the greenhouse under a 12 h light period at 24 °C and a 12 h dark period at 16 °C, with 60% relative humidity (Salih et al., 2024). Regular watering kept the soil mix moist, and plants were fertilized with NPK (20:20:20) as needed.

### Plasmodiophora brassicae inoculation

One milliliter of 1.0 × 10^8^ spores/mL resting spore suspension of P. brassicae pathotype 3H (Canadian clubroot differential set) and AB_B_2024-22 isolates was inoculated onto seven-day-old canola seedlings through the soil around the roots. At 42 days post-inoculation (dpi), plant roots were harvested and evaluated for clubroot symptoms.

### DNA extraction, resistance gene enrichment, and genome sequencing

High molecular weight DNA samples were extracted from three-week-old canola leaf samples for RenSeq and whole genome sequencing. DNA samples were extracted by an optimized CTAB protocol (Gonzalez-Garcia et al., 2026). The quality and quantity of DNA were determined by Nanodrop oneC and Qubit 4 (Thermo Fisher Scientific, MA, USA), respectively. For the RenSeq procedure, the NLR bait library was designed based on the “ZS11” B. napus NLR gene sequences (Chen et al., 2021), and the capture of the NLR enrichment library was carried out by Arbor Bioscience (Daicel Arbor Biosciences, Ann Arbor, MI) using the myBaits Hyb Capture Kits standard protocol version 5.02 (https://arborbiosci.com/wp-content/uploads/2022/03/myBaits_v5.02_Manual.pdf).

Afterward, these target-captured libraries were sequenced on the Illumina NovaSeq 6000 platform (Illumina Inc., San Diego, CA). Paired-end libraries with an insert size of 150 bp reads. After the adaptor trimming, the sequenced reads were processed using FastQC for quality control. Genome sequencing was performed using the Oxford Nanopore Technology (ONT) MinION platform (R9.4.1 flow cell) with ∼10x coverage, and Guppy v6.4.6 was used for base calling. ONT raw sequencing reads were filtered to remove the unreliable data (Q Score < 7, fragment < 1kb) using NanoFilt (De Coster et al., 2018). All passed reads were applied for further analysis.

### Genome Assembly and Annotation

The filtered ONT reads were assembled using Flye v2.9 (Kolmogorov et al., 2019). The Flye release was set to a genome size of 930m. After that, we applied Medaka v1.8.0 for the first-round polishing of the assembled data using the parameter “-m r941_min_high”. The raw RenSeq reads were used for the second round of genome correction by Hapo-G v1.2 (Aury & Istace, 2021). After two rounds of error correction, we finally obtained the contig-level draft genomes of five canola lines.

Obtained contigs were filtered to >5kb and used to predict genes in the next step. The genes from the five polished assembled B. napus genomes were annotated ab initio using Helixer v0.3.3 (https://www.plabipd.de/helixer_main.html) with the land plant model (land_plant_v0.3_a_0080) (Holst et al., 2025). Then, we extracted protein sequences using AGAT v1.2.0 (Dainat et al., 2023) from the GFF3 files generated by Helixer.

### Identification of NLR genes and phylogenetic analysis

Extracted protein sequences were used for NLRtracker to identify NLR genes (Kourelis et al., 2021). All these NLRs were classified into eight subgroups: CC-NBARC, CC-NLR, NBARC, NBARC-LRR, RPW8-NBARC, RPW8-NLR, TIR-NBARC, and TIR-NLR. To identify the NLR-ID, we used InterProScan v5 (Jones et al., 2014) to search for integrated domains in NLR amino acid sequences using the Pfam, Gene3D, and SUPERFAMILY databases, with an e-value cutoff of < 10^-5^.

To investigate the phylogenetic relationship of NLRs of five canola lines. The conserved NBARC domain of NLR protein sequences was extracted to construct the phylogenetic tree of the NLRs. These extracted sequences were aligned and trimmed using MUSCLE v5.1 (Edgar, 2022) and trimAI v1.5.1 (Capella-Gutiérrez et al., 2009), and the resulting files can be used as input for the analysis. The maximum-likelihood NBARC domain phylogeny of 2,278 amino acid sequences was inferred using IQ-TREE 2 (Minh et al., 2020) with the JTT+F+R9 model and 1,000 bootstrap replicates. Color blocks represent different types of NLRs, including CC-NBARC, CC-NLR, RPW8-NBARC, RPW8-NLR, TIR-NBARC, TIR-NLR, NBARC, and NBARC-LRR.

### Pairwise reciprocal best hits (RBH) and Orthogroups (OGs) identification

For the pairwise reciprocal best hits (RBH) analysis, we ran MMseqs2 easy-rbh on every pair of the six per-line FASTAs with the “-e 1e-10 --min-seq-id 0.4 -c 0.8 --cov-mode 0” parameters (Kallenborn et al., 2025). The B. napus pan-NLRome was constructed using the protein clustering approach. Firstly, we performed all-against-all comparisons of full-length NLR proteins from five CR lines and the Westar line using DIAMOND (Buchfink et al., 2015) inside the OrthoFinder v2.5.5 software (Emms & Kelly, 2019) and grouped them based on orthology. The pan-NLRome was classified into four categories: core (present in all six lines), soft core (present in five lines), shell (present in two to four lines), and private (present in one line). For the phylogenetic analysis of the NLR OGs, we extracted the NBARC domain from each OG and used IQ-TREE 2 (Minh et al., 2020) to build the maximum-likelihood tree with the JTT+F+R9 model and 1,000 bootstrap replicates.

### Synteny analysis

The CRA 3.7 and CRA 8.2 regions were retrieved from the ECD04 genome (Yang et al., 2022), and the contigs containing the candidate regions of CRa and Crr1a were retrieved from each assembly. Synteny analysis between the reference genome ECD04 and each IH line was performed using MCscan (Python version).

### Amplification of the clubroot resistance CRa and Crr1a gene

The extracted HMW DNA samples from Westar and five CR lines were used to amplify the CRa and Crr1a genes to assess allelic variation. Phusion™ High-Fidelity DNA Polymerase (Thermo Fisher Scientific, MA, USA) was used for gene amplification, and PCR was carried out in 50 μL reaction mixtures according to the manufacturer’s instructions. The PCR primer pairs for amplification are listed in the Supplementary Table S1. The amplicons were sent to the Plasmidsaurus (South San Francisco, CA, USA) for linear amplicon sequencing.

### Predication and visualization of protein structures

AlphaFold3 (Abramson et al., 2024) was used to predict protein structures in this study. Then, we utilized UCSF ChimeraX 1.10.1 for structural visualization and alignment (Meng et al., 2023). The protein structural comparison analysis was conducted by TM-align software (Version 20220412) (Zhang & Skolnick, 2005).

### Data analysis and visualization

The UpSet plot and the chromosome physical map were generated by R using the UpSetR package (Conway et al., 2017) and the ggplot2 package (Villanueva & Chen, 2019), respectively. We used IBS 2.0 to illustrate protein domain structures (Xie et al., 2022). Phylogenetic trees were visualized using the tvBOT (Xie et al., 2023) web platform (https://www.chiplot.online/tvbot.html).

## ACKNOWLEDGEMENTS

The authors thank the Daicel Arbor Biosciences team for their guidance during bait design and data analysis. We are also thankful to Fonds de recherche du Québec - Nature et technologies (Grant ID: 350513) for providing the doctoral scholarship to JW.

## AUTHOR CONTRIBUTIONS

EPL conceived and funded the project. EPL, JW, and MS planned and designed the research. JW, YA, MS, CF, and EPL contributed to the methodology. JW, MAJ, EF, and EPL performed experiments and formal analysis. CF provided the seed resource. JW and EPL prepared the figures and tables. JW and EPL wrote the original draft of the manuscript. All authors contributed to the review and editing of the manuscript.

## DATA AVAILABILITY

The raw data generated in this study are available in the Bioproject PRJNA1459298. The code used can be found in the GitHub link: https://github.com/Edelab/CR_pan_NLRome. The rest of the data that supports the findings are available in the Supplementary materials.

## SUPPLEMENTARY DATA

**Supplementary Table S1.** Primers used for PCR analyses.

**Supplementary Table S2.** Resistant profiles of 5 CR B. napus lines

**Supplementary Table S3.** NLR numbers by each class in 5 CR B. napus lines

**Supplementary Table S4.** Full list of NLRs with C-JID domains of 5 CR lines generated by NLRtracker

**Supplementary Table S5.** Full list of NLRs with integrated domains of 5 CR lines

**Supplementary Table S6.** pairwise reciprocal best hits (RBH) of six lines

**Supplementary Table S7.** pan-NLRome orthogroups and the NLR names

**Supplementary Table S8.** Pan-NLRome of NLR copy numbers of each line

**Supplementary Table S9.** Pan-NLRome of presence/absence and classification

**Supplementary Table S10.** The Westar NLR of the core OGs location on 19 chromosomes

**Supplementary Table S11.** The candidate orthogroups based on different CR profiles

**Supplementary Table S12.** The 31-candidate resistance-enriched orthogroups

**FIGURE S1.**
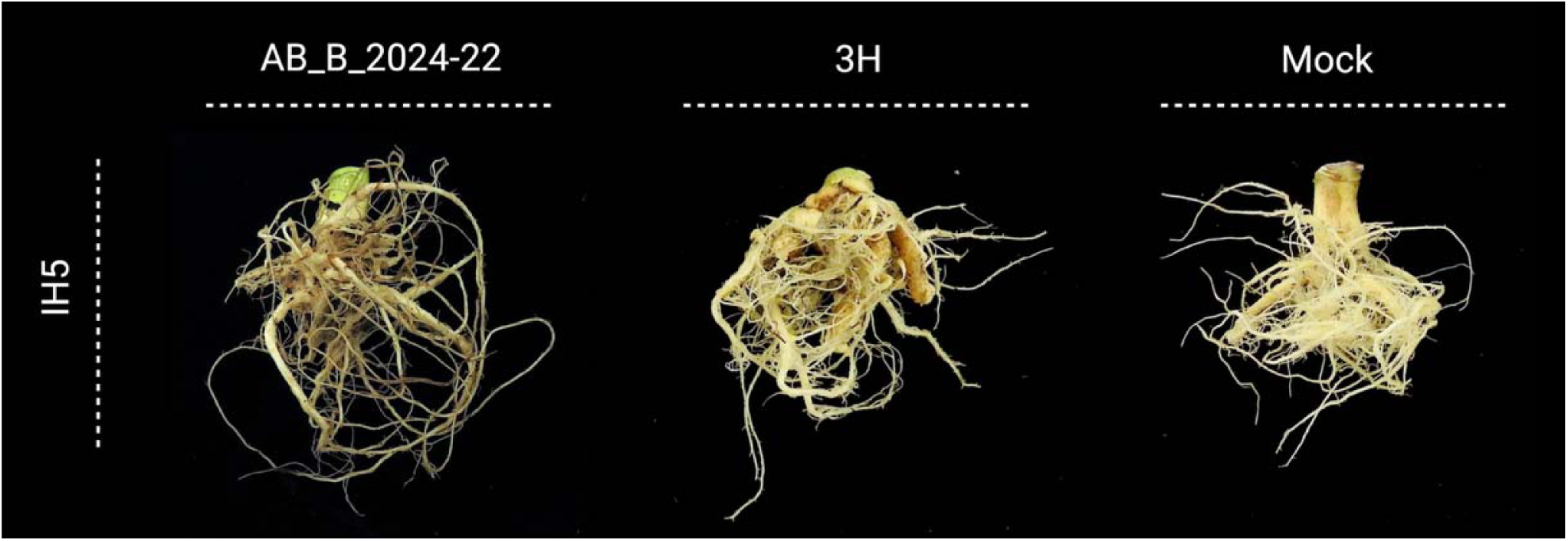
Representative root phenotypes of CR line IH5 following inoculation with P. brassicae pathotype AB_B_2024-22 (highly virulent) and 3H (less virulent), alongside mock-treated controls.

**FIGURE S2.**
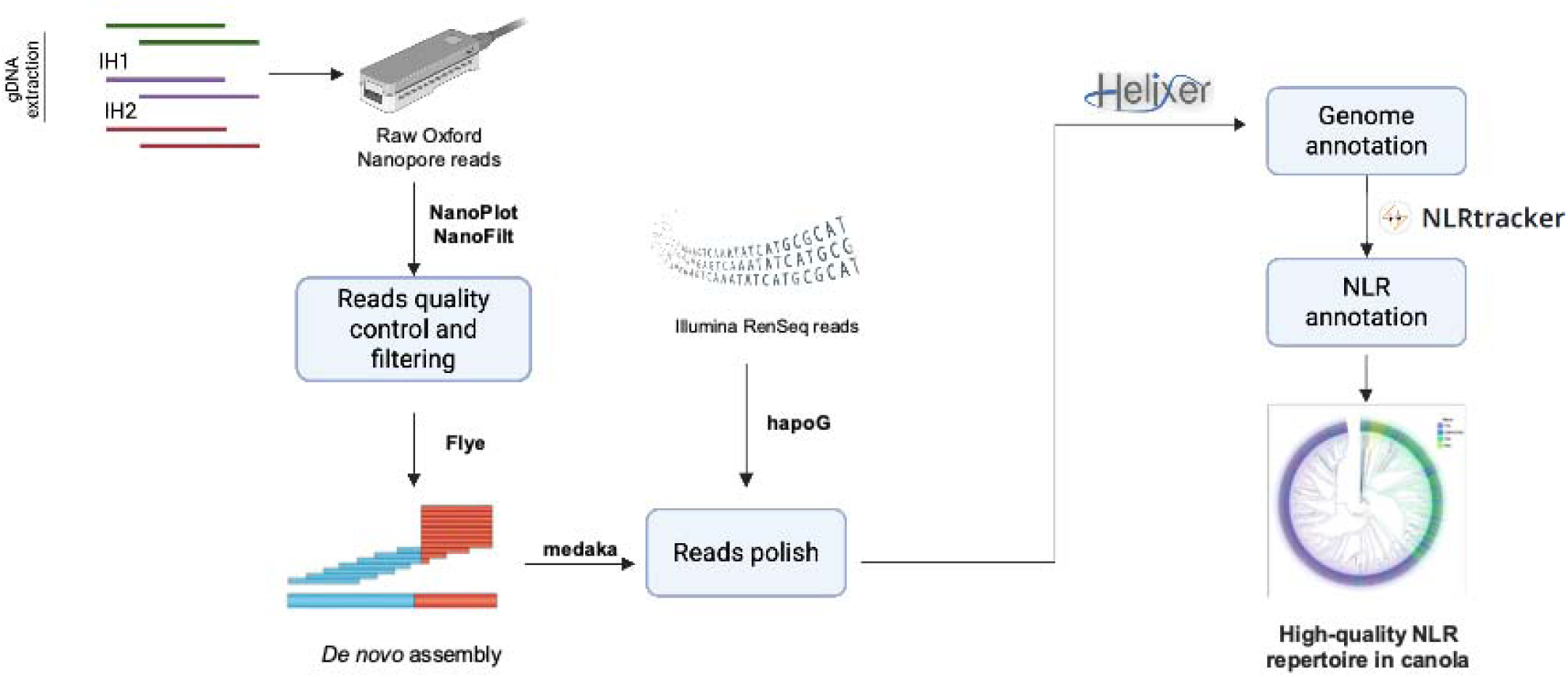
Workflow and pipeline for constructing a high-quality NLR repertoire in CR canola. Raw ONT reads were first processed using NanoPlot and NanoFilt for read quality assessment and filtering. The filtered long reads were assembled de novo using Flye, followed by the first round of polishing with Medaka. After that, Hapo-G was used for the second-round polishing using RenSeq reads from these five lines. The polished genome assembly sequences were then used to annotate the genome with Helixer. Predicted gene models were further screened using NLRtracker to identify and classify NLR genes. This integrated long-read assembly, RenSeq-guided refinement, genome annotation, and NLR classification pipeline generated a high-quality NLR repertoire for five CR canola lines.

**FIGURE S3.**
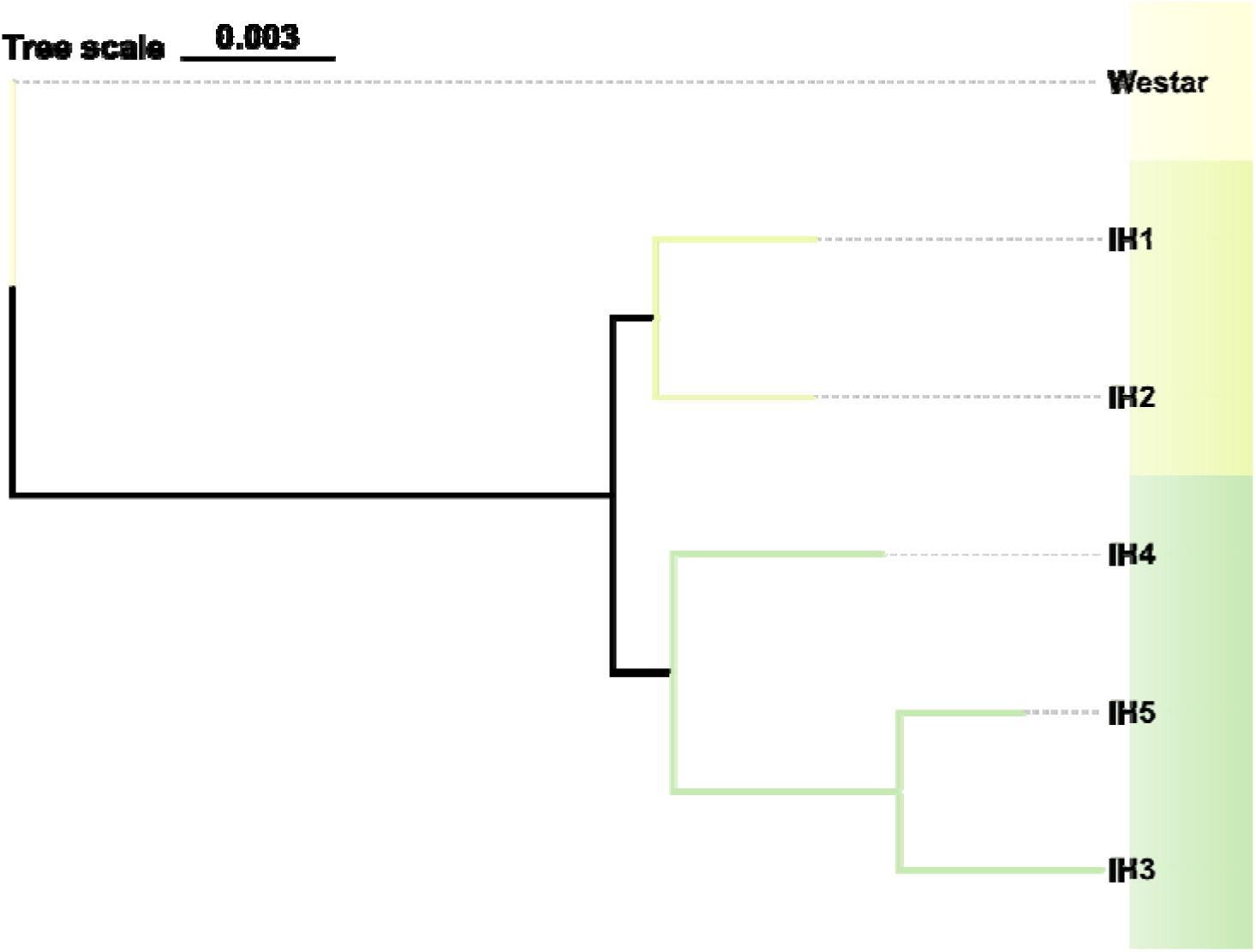
Species tree of the six B. napus lines based on single-copy NLR orthologs. Tree scale is 0.003.

**FIGURE S4.**
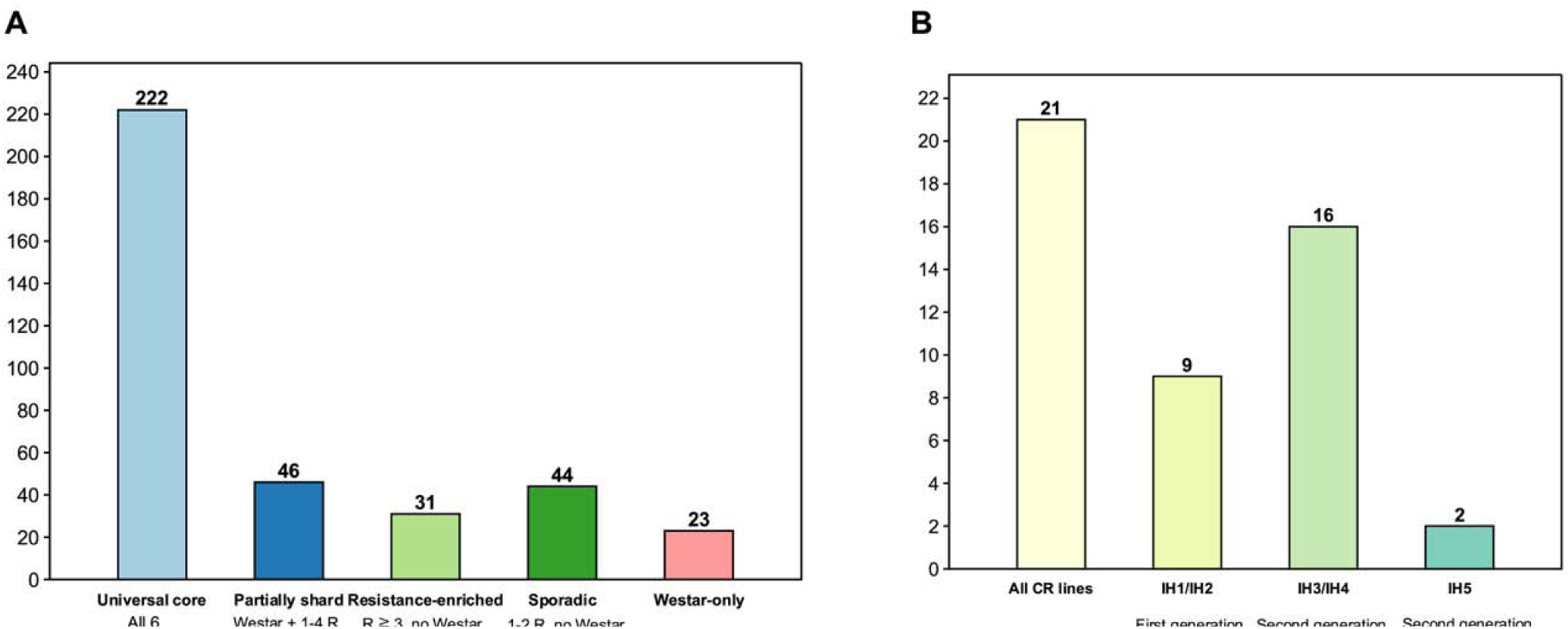
Pan-NLRome orthogroup partitioning across the six canola lines. **(A)** Pan-NLRome categories based on co-occurrence across the six lines. All 366 NLR OGs were assigned to one of five mutually exclusive categories. (**B**) OGs partitioned by different CR types of the contributing lines.

**Figure S5.**
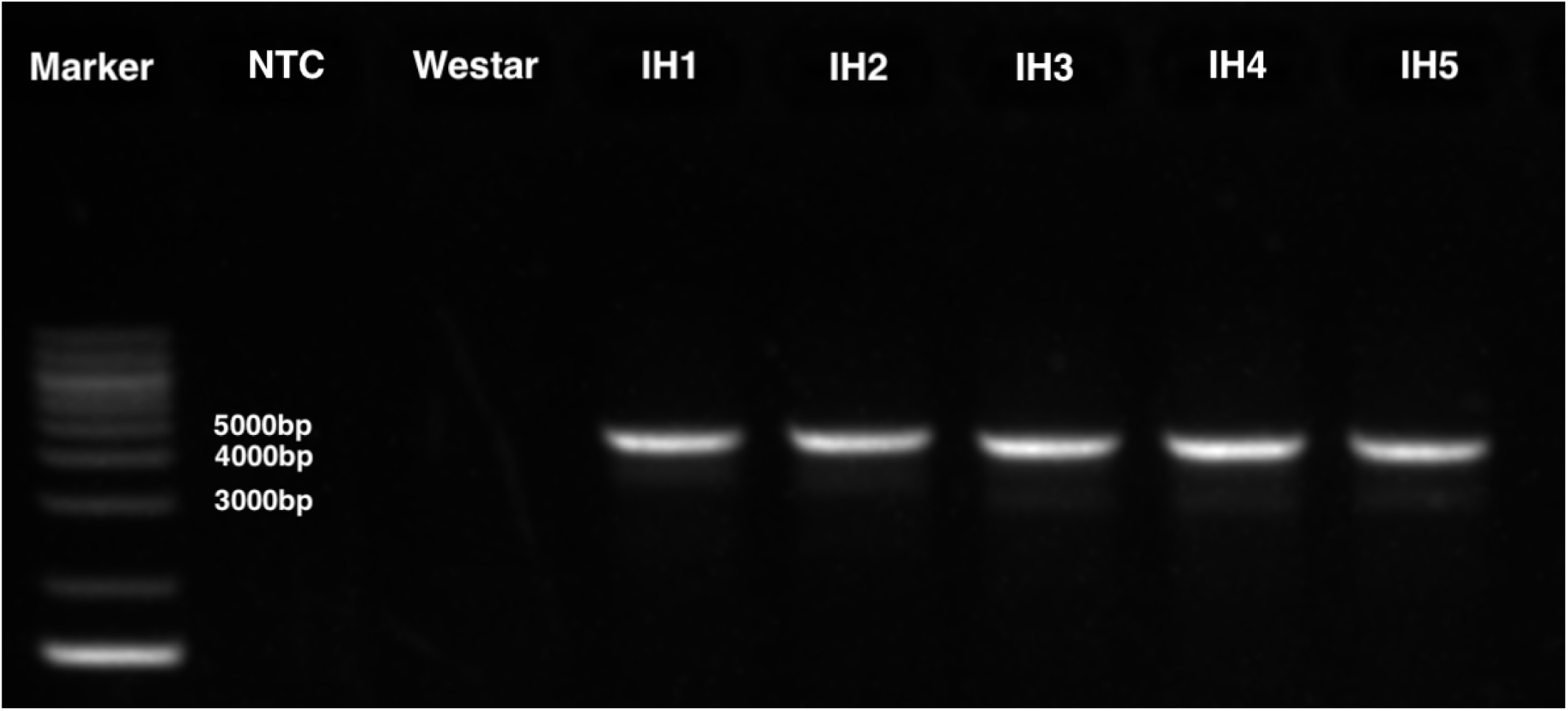
CR amplification of the CRa gene in five clubroot-resistant canola lines. Agarose gel electrophoresis showing PCR amplification of the CRa gene in Westar and five CR canola lines (IH1-IH5). A clear amplicon of approximately 4-5 kb was detected in all five CR lines, IH1-IH5, whereas no visible amplification was observed in the no-template control (NTC) or the susceptible cultivar Westar. The DNA marker is shown in the left lane. These results support the presence of CRa-like sequences in the resistant lines.

